# A Feedback Motif that Breaks the Fundamental Limit on Noise Suppression to Stabilize Fate

**DOI:** 10.1101/141390

**Authors:** Winnie Y. Wen, Elena Ingerman, Maike M. K. Hansen, Brandon S. Razooky, Roy D. Dar, Charles Chin, Michael Simpson, Leor S. Weinberger

**Affiliations:** Gladstone Institutes (Virology and Immunology), San Francisco, CA 94158 United States; Bioinformatics and Systems Biology Graduate Program, University of California, San Diego, La Jolla, CA 92093 United States; Center for Nanophase Materials Science, Oak Ridge National Laboratory, Oak Ridge, TN 37831 United States; Department of Biochemistry and Biophysics, University of California, San Francisco, CA 94158 United States

**Keywords:** stochastic noise, fate selection, negative feedback, post-transcriptional splicing

## Abstract

Diverse biological systems utilize gene-expression fluctuations (‘noise’) to drive lineage-commitment decisions^1-5^. However, once a commitment is made, noise becomes detrimental to reliable function^6,7^ and the mechanisms enabling post-commitment noise suppression are unclear. We used time-lapse imaging and mathematical modeling, and found that, after a noise-driven event, human immunodeficiency virus (HIV) strongly attenuated expression noise through a non-transcriptional negative-feedback circuit. Feedback is established by serial generation of RNAs from post-transcriptional splicing, creating a precursor-product relationship where proteins generated from spliced mRNAs auto-deplete their own precursor un-spliced mRNAs. Strikingly, precursor auto-depletion overcomes the theoretical limits on conventional noise suppression—minimizing noise far better than transcriptional auto-repression—and dramatically stabilizes commitment to the active-replication state. This auto-depletion feedback motif may efficiently suppress noise in other systems ranging from detained introns to non-sense mediated decay.

---

Stochastic fluctuations (noise) in gene expression are inescapable at the single-cell level^8-11^. These fluctuations can be beneficial, enhancing fitness in variable environments^1,2^, but are also detrimental to reliable cellular function and subject to purifying evolutionary selection^6,7^. For example, during lineage-commitment decisions, transcriptional noise promotes selection between alternate cell fates^12-14^ but, after commitment, such noise is presumably attenuated^15,16^ to enable reliable cell function. The mechanisms enabling mammalian cells to transition from noise-enhancing to noise-suppressing states remain unclear, and the putative mechanisms are subject to known limits on noise suppression^17^.

The life cycle of Human Immunodeficiency Virus Type 1 (HIV-1, hereafter HIV) encodes a stereotyped fate-commitment decision^18^ where the virus undergoes a noise-driven binary fate decision (Fig. 1A) leading to either an active replication program—producing viral progeny and cell death—or to a long-lived quiescent state called proviral latency^19,20^. The viral fate decision is driven by large stochastic fluctuations in the activity of HIV’s long terminal repeat (LTR) promoter, which are amplified by positive feedback from HIV’s Tat protein and are sufficient to shift cells between active replication and latency^12,21^. Whether these large transcriptional fluctuations are attenuated after commitment to active replication is not known.

**Figure 1:**
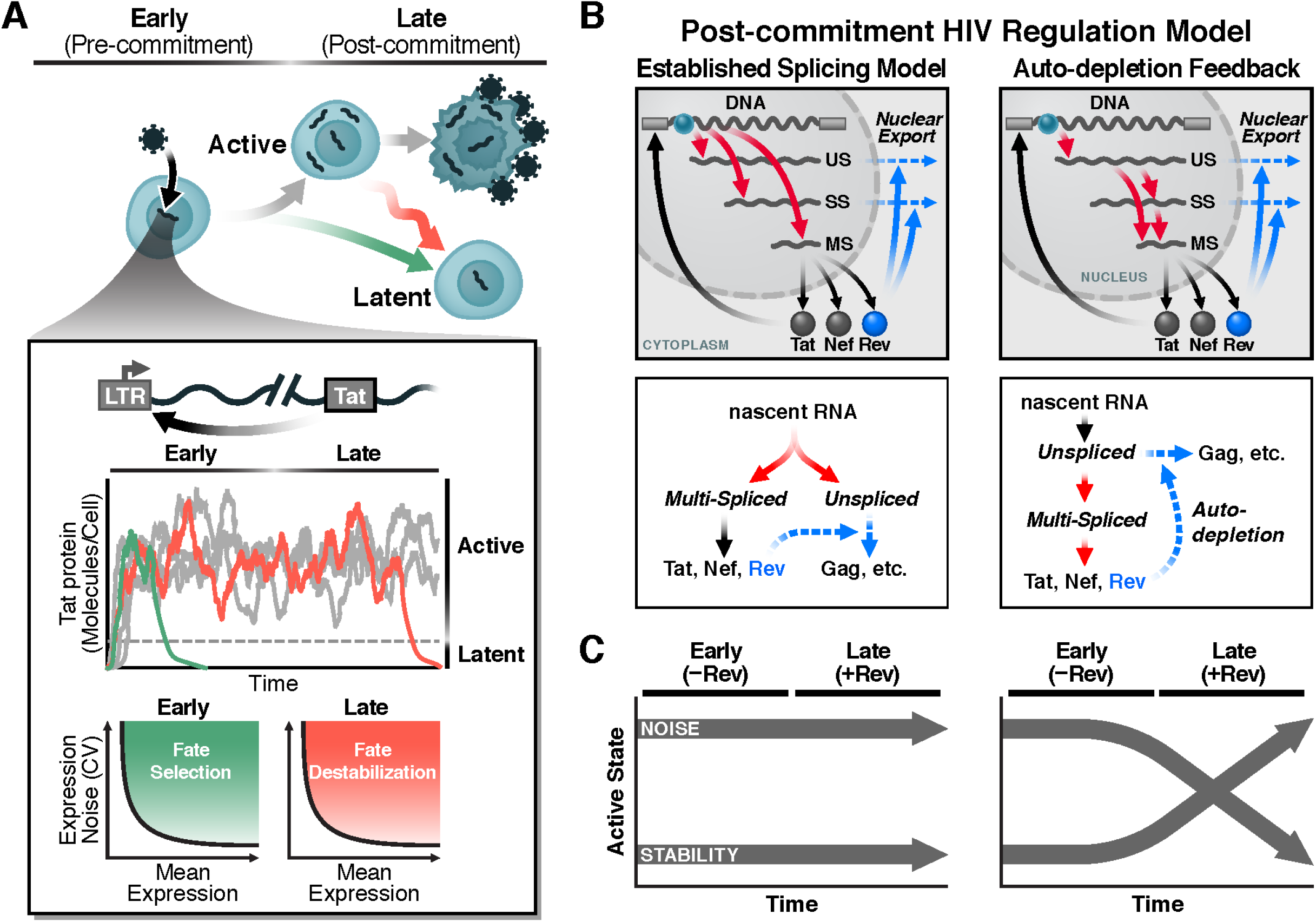
Fate stabilization in noise-driven fate-selection circuits. **(A)** HIV’s fate decision in CD4^+^ T cells: early after infection, infected cells commit to either latent or active infection with actively infected cells producing virus after a delay of about 1 day and then dying after about 2 days. Inset: the HIV Tat positive-feedback circuit regulates the active and latent states with Tat transactivation of the LTR promoter (HIV’s only promoter) being obligate for active replication and low Tat levels enabling viral latency. As long as positive feedback persists, it generates large fluctuations in gene expression, enabling entry to latency (green) but potentially destabilizing the active state (red) where black line is predicted noise-mean relationship from a Poisson model. (**B**) Two models of HIV gene regulation during active replication. Left: The established model of co-transcriptional RNA alternative splicing; 9-kb unspliced (US), 4-kb singly spliced (SS), and 2-kb multiply spliced (MS) RNAs are independently generated and thus Rev-mediated export of US and SS does not deplete MS RNA. Right: The precursor auto-depletion model relies on post-transcriptional splicing where US and/or SS RNAs are precursors for MS RNA and Rev-mediated export of US and SS precursors depletes MS RNA establishing indirect negative feedback. (**C**) Differing noise and stability properties between the two models: both models enable high noise and fate selection pre-Rev expression (early) but precursor auto-depletion allows attenuation of noise and stabilization of the active state upon Rev expression (late).

HIV encodes components of a putative negative-feedback loop^22^, which could attenuate noise ^23^. However, models of co-transcriptional splicing in HIV^24,25^ are inconsistent with this reported negative-feedback mechanism, which requires a precursor-product relationship and post-transcriptional splicing (Fig. 1B). Specifically, HIV pre-mRNA is spliced into three classes of transcripts: a 9-kb un-spliced (US) transcript, a 4-kb singly spliced (SS) class, and a 2-kb multiply spliced (MS) class. The US and SS transcripts (encoding capsid, envelope, and genomic RNA) are initially retained in the nucleus of infected cells, but the MS transcripts, which encode the regulatory proteins Tat, Rev, and Nef, are efficiently exported. Nuclear export of the US and SS transcripts is facilitated by binding of Rev proteins to the Rev-Responsive Element (RRE) within the US and SS transcripts, which enables export through cellular chromosome region maintenance 1 (CRM1)^26-28^. The essential consequence of the co-transcriptional model is that a transcript’s splicing fate is sealed during transcriptional elongation, such that differentially spliced transcripts are uncoupled from each other and share no precursor-product relationship (Fig. 1B, left). Thus, upon Rev-mediated export of US RNA, the co-transcriptional splicing model precludes any effect on MS RNA (i.e., no negative feedback), enabling the persistence of destabilizing stochastic fluctuations (Fig. 1C, left). Conversely, HIV splicing occurring post-transcriptionally would establish a precursor-product relationship and enable negative feedback upon Rev-mediated export of US RNA (i.e., precursor auto-depletion), thereby attenuating transcriptional noise and stabilizing commitment to the active-replication state (Fig. 1, B and C, right).

We used single-molecule mRNA Florescence *In Situ* Hybribidation (smFISH) (Fig. 1A) to examine whether HIV splicing is co-transcriptional^24,25^ or post-transcriptional^29,30^. To discriminate between co- and post-transcriptional splicing, we performed a transcriptional pulse-chase experiRef26ment^31^ where HIV transcription was activated by tumor necrosis factor alpha (TNF) and chased 14 minutes later by promoter shutoff using the transcriptional elongation inhibitor actinomycin D (ActD) (Fig. 1, B and C). Cumulatively, more than 2000 cells were analyzed by smFISH for this experiment (about 100 cells per time point). During the 14-minute TNF pulse, US RNA substantially increases (whereas neither SS nor MS RNA increases), whereas during the ActD chase, US RNA is depleted while SS RNA “peakes” (first increases and then decreases) and MS RNA continually increases. These results indicate that SS and MS RNA are spliced post-transcriptionally from a US RNA precursor, since both species increase after transcriptional elongation is halted. An alternative spatial assay of post-transcriptional splicing^32^—which quantifies the radial distance of individual mRNA molecules from the transcriptional center— was also inconsistent with co-transcriptional splicing showing that SS and MS mRNAs are maximal ~1.5 and 3 μm from the transcriptional center, respectively, and only US mRNAs were observed at transcriptional centers (Fig. S1, C and D). As expected from Stokes-Einstein diffusion theory, temporal distributions of splicing (Fig. S1E) agree with these spatial distributions. To further test whether transcription is dispensable for splicing, we developed an assay termed <u>S</u>plicing <u>A</u>fter <u>T</u>ransfection of <u>U</u>nspliced pre-m<u>R</u>NA into the <u>N</u>ucleus (SATURN) in which synthetic, *in vitro* transcribed full-length HIV mRNA was transfected into cellular nuclei. This assay relies upon a reporter expressed only from a multiply-spliced transcript and tests whether it is possible for splicing to occur, without *in situ* transcription within cells. Over half of transfected T cells exhibited MS reporter expression 4 hours after transfection of full-length unspliced HIV mRNA (Fig. S2), demonstrating that HIV can be spliced post-transcriptionally within cells. Overall, the results establish that MS RNA is serially generated from US RNA (or SS RNA) leading to a precursor-product relationship—the necessary criterion for precursor auto-depletion feedback.

To determine whether HIV-expression dynamics exhibit negative feedback, we next used quantitative time-lapse microscopy. Mathematical models (Supp. Info) showed that co-transcriptional splicing generates the expected monotonic increase to steady state, whereas post-transcriptional splicing generates precursor-auto depletion and leads to ‘overshoot’ kinetics characteristic of negative feedback (Fig. 2D). To examine the kinetics of HIV gene expression in single cells, we stably transduced cells with an inducible HIV-reporter virus. The reporter virus is a single-round (Δenv) HIV^33^ encoding a short-lived green fluorescent protein with a two-hour half-life (d_2_GFP) within the HIV Nef reading frame—each transduced cell contains a single, latent integration of HIV d_2_GFP that is inducible by TNF and the Δenv mutation ensures no subsequent transmission of the virus between cells after treatment with TNF. Longitudinal single-cell imaging over 25 hours after TNF treatment shows that virtually all cells exhibit GFP increasing to reach maximal abundance at 5 to 8 hours, followed by a decay in GFP intensity (Fig. 2E and Movie S1), as predicted by negative feedback. Moreover, a subset of cells exhibited multiple, damped oscillations in GFP (Fig. S3)—another hallmark of negative feedback^34,35^—and temporal auto-correlation of the single-cell trajectories, a sensitive reporter of feedback streRef37ngth^36,37,^ further confirmed negative feedback (Fig. S4). Similar overshoot trajectories were observed after direct infection of human, donor-derived primary CD4^+^ T cells (Fig. S5) and primary, monocyte-derived macrophage cells (Fig. S6), indicating that negative feedback operates during infection of primary cells and latent reactivation.

**Figure 2:**
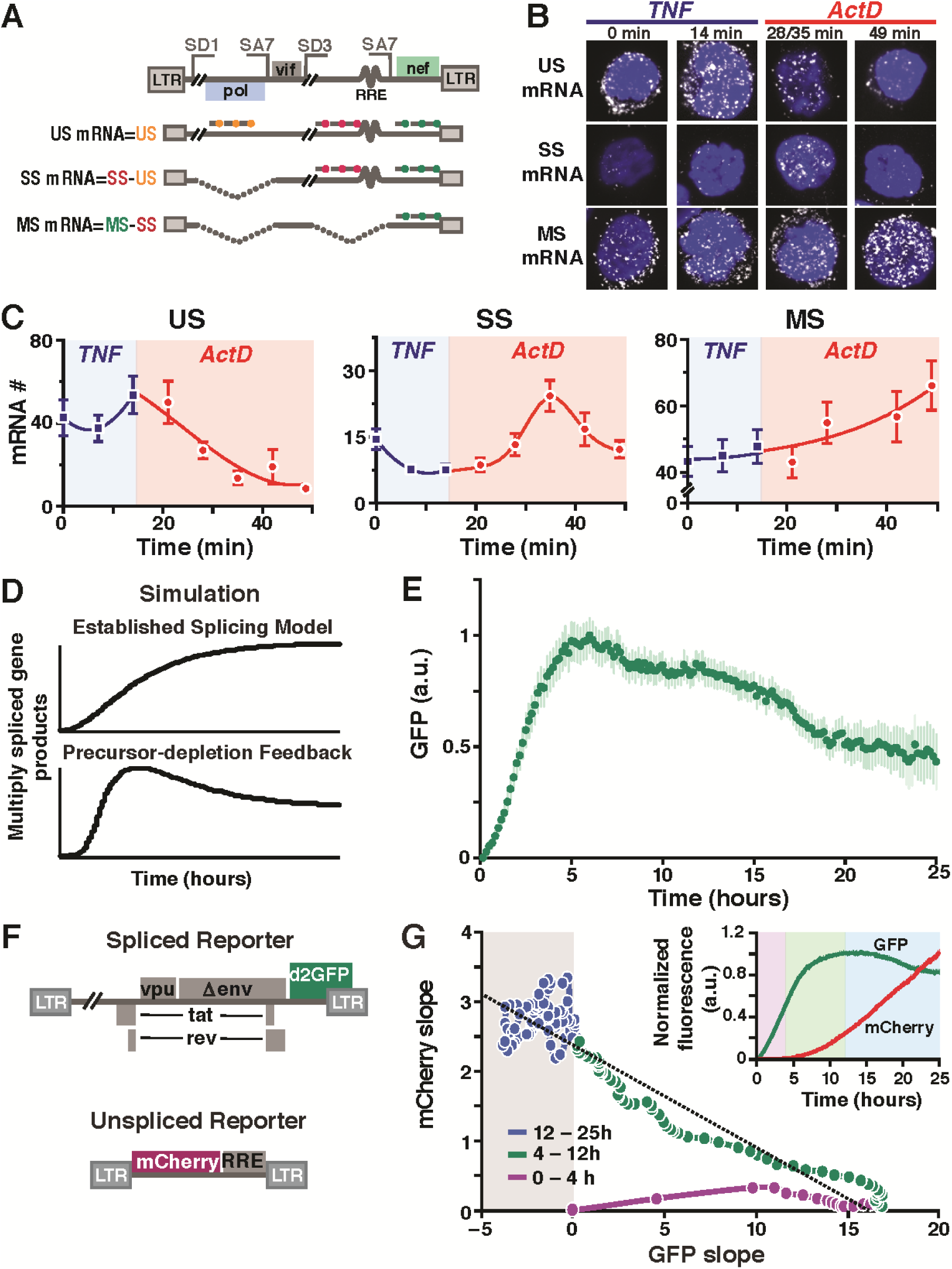
HIV mRNAs are spliced post-transcriptionally, establishing a precursor-product relationship and enabling precursor auto-depletion feedback. **(A)** Schematic of smFISH probes used to quantify US, SS and MS mRNAs shown relative to the HIV-1 genome with representative splice donor (SD) and splice acceptor (SA) sites and representative US, SS, and MS HIV genes. (**B**) Representative smFISH images of transcriptional pulse-chase in HIV-infected Jurkat cells. TNF, activation of the HIV promoter was chased 14 minutes later with the transcriptional elongation inhibitor Actinomycin D (ActD). (**C**) Quantification of mRNA molecules during the TNF pulse (blue) and the ActD chase (red). Each data point is the mean nuclear mRNA count for about 100 cells with trend lines and error bars represent SEM. (**D**) ODE-model trajectories showing disparate predictions for co-transcriptional and post-transcriptional splicing models of HIV (see Supp. Info for equations). (**E)** Time-lapse imaging of Jurkat cells transduced with HIV d_2_G after induction with TNF (cells are monoclonal for the HIV-integration site). (**F**) Schematic of two-color reporter system showing the MS reporter (d2GFP) in *nef* reading frame and the US reporter (mCherry) being dependent on Rev-mediated export. (**G**) Direct coupling of negative feedback to Rev-mediated export of unspliced RNA export. Quantification of GFP and mCherry slopes: (i) during the early phase after TNF stimulation (0 to 4 hours), GFP production rate is positive and increasing while mCherry production is near zero, consistent with Rev needing to accumulate and multimerize for its function ^52^; (ii) as mCherry production increases (4 to 12 hours), GFP production begins to taper off, consistent with depletion of MS RNA; (iii) in the late phase (after 12 hours), the GFP production rate is negative only when the mCherry production rate is high. Inset: mean GFP and mCherry trajectories of 100 representative cells.

To verify that the overshoot dynamics were not caused by transcriptional silencing or TNF-induced oscillatory dynamics^35^, we performed four different tests. First, we tested an array of HIV-reporter constructs lacking Rev; all of these constructs generate monotonic increases in GFP, rather than overshoots, in cells treated with TNF (Fig. S7). We tested whether subsequent re-stimulation of cells with TNF, during the overshoot, or subsequent re-stimulation of cells with the histone deacetylase inhibitor trichostatin A, during the overshoot, would re-stimulate the HIV LTR and abrogate the overshoot, as transcriptional silencing predicts^38^; no abrogation of the overshoot occurs upon re-stimulation (Fig. S8). To confirm that the precursor-product relationship is responsible for the overshoot, we tested whether strengthening Rev export would enhance auto-depletion and intensify the overshoot and found that a construct encoding a second distal RRE (in *gag*) exhibits faster overshoot-decay kinetics than HIV-d_2_GFP (Fig. S9). Leptomycin B, an inhibitor of Rev export^39^, diminishes the overshoot, (Fig. S10), indicating Rev export and not transcriptional silencing, mediates the overshoot.

To determine whether the observed overshoot is directly coupled to Rev export of unspliced RNA precursors, we developed a two-color reporter system (Fig. 2F) in which a second construct, LTR-mCherry-RRE, requires Rev nuclear export for mCherry expression, and acts as a Rev-dependent reporter for US RNA expression^40^. As in the single-color reporter system, GFP is produced from a multiply-spliced transcript and provides a measure of MS RNAs. Analysis of GFP and mCherry intensities in individual cells (Fig. 2G, inset) shows that thRef8e GFP overshoot occurrs concurrently with increasing expression from the US mCherry reporter (the reduced GFP overshoot relative to the one-color system is consistent with Rev having to export an extra species of US mCherry RNA). Plotting the relative slopes (i.e., net-expression rates^8^) verifies that expression from US transcripts precisely coincides with the decay in expression from MS transcripts (Fig. 2G). This coincident reversal in slope was confirmed by smFISH (Fig. S1D). Together, these data demonstrate that Rev-mediated nuclear export of its own unspliced precursor RNA underlies this negative-feedback circuitry.

We next asked if this post-transcriptional form of negative feedback (precursor auto-depletion) could provide any theoretical benefit over archetypal transcriptional auto-repression. Computational and analytical arguments (Box 1 and Supp. Info) indicate that negative feedback through auto-depletion of precursor RNA attenuates noise more effectively than transcriptional auto-repression. The improved noise attenuation is predicated on splicing being inefficient compared to export, which is also a necessary criterion for auto-depletion feedback. A model of HIV gene regulation was developed and extensively validated (Supp. Info.) and this model also predicts that enhancing the efficiency of HIV splicing diminishes precursor auto-depletion (Fig. 3A). To experimentally test this modeling prediction, we focused on the HIV splice acceptor SA7, which is used by all three MS transcripts (Tat, Rev, and Nef) but lacks the canonical sequence elements required for efficient splicing; i.e., SA7 exhibits neither a consensus branch-point A nor a poly-pyrimidine tract^41^. A series of SA7 HIV mutants was generated to enhance the poly-pyrimidine tract and create a consensus branch-point A sequence (Fig. S11A). These optimized SA7-mutant viruses exhibit enhanced RNA splicing (Fig. S12) and increased GFP expression, but lack the overshoot kinetics characteristic of negative feedback (Fig. 3B). Despite an increase in mean-expression of GFP, the optimized SA7 mutants also show increased noise (Fig. 3, B and C) and substantial decoupling between MS and US expression kinetics by two-color analysis (Fig. 3D). This noise increase, although contrary to simple Poissonian models—in which noise decreases as the mean-expression level increases—is consistent with diminution of negative feedback, which suppresses both noise and mean-expression level. The data indicate that inefficient splicing is an important molecular determinant of precursor auto-depletion and its resultant noise minimization.

**Figure 3:**
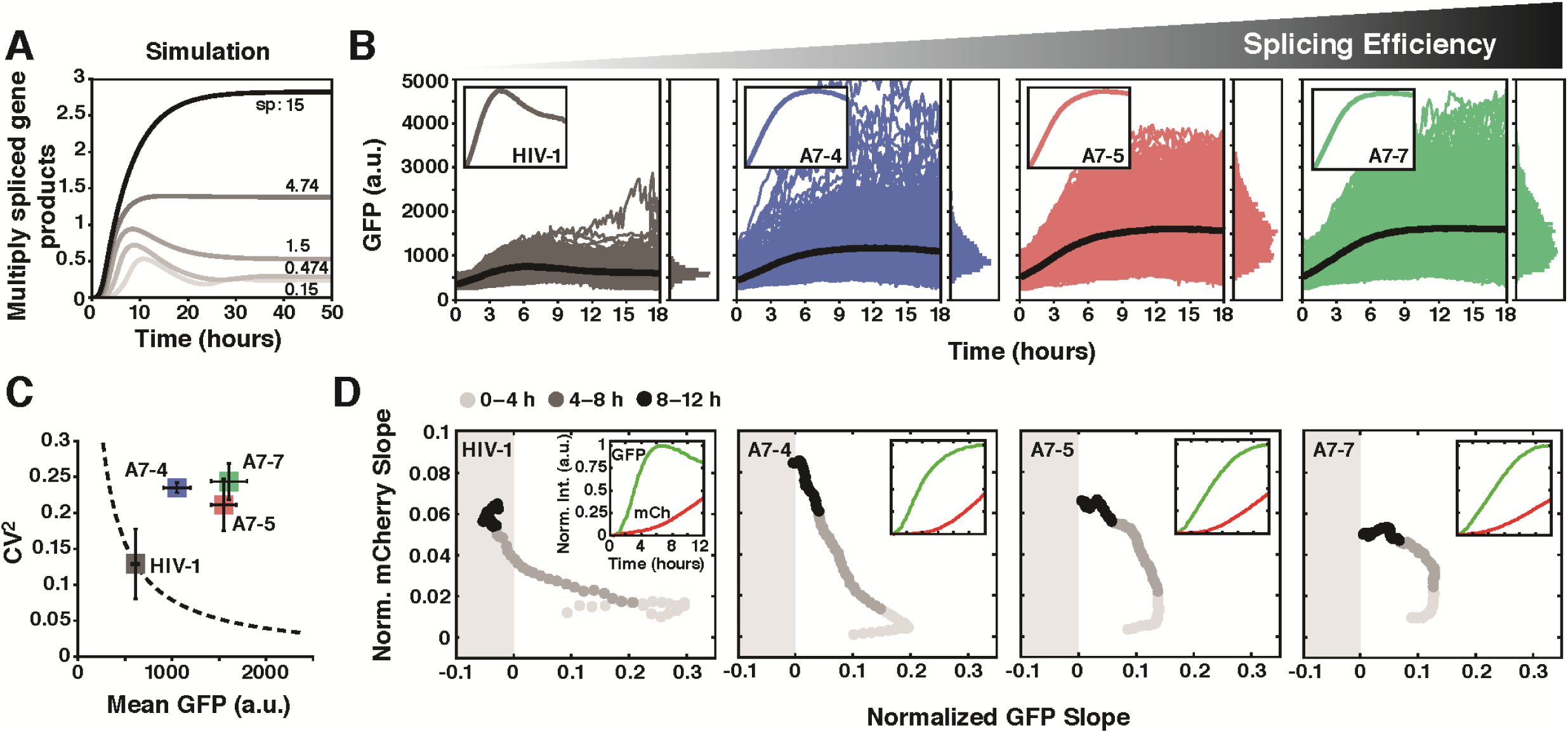
Precursor auto-depletion reduces HIV expression noise. **(A)** Output of HIV gene-regulation computational model validated on single-cell trajectory data (Supp. Info, see Table S10). Model predicts that enhancement of splicing abrogates precursor auto-depletion feedback. (**B**) Time-lapse microscopy of wild-type HIV d_2_G and three SA7 mutants with enhanced splicing (populations are polyclonal for viral integration sites). Insets: mean trajectories normalized to max (to examine overshoot). (**C**) CV^2^ versus mean for data in panel B shows that mutants exhibit large increase in CV^2^ despite increased mean expression. Dotted line is the Poisson-model prediction. (**D**) Decoupling between MS and US expression in the SA7 mutants. The two-color reporter assay shows that, in the SA7 mutants, US expression rate (i.e., mCherry slope) increases without the wild-type-like decrease in MS expression rate (i.e., d_2_GFP slope) and MS expression rate does not enter the negative (i.e., negative feedback) regime. Slopes are normalized to max for comparison purposes. Insets: mean intensity trajectories for GFP and mCherry.

### Box 1: Breaking the Limit on Noise Suppression

Noise in gene expression is often analyzed (Alon, 2007), by comparing standard two-state models of gene expression as follows:

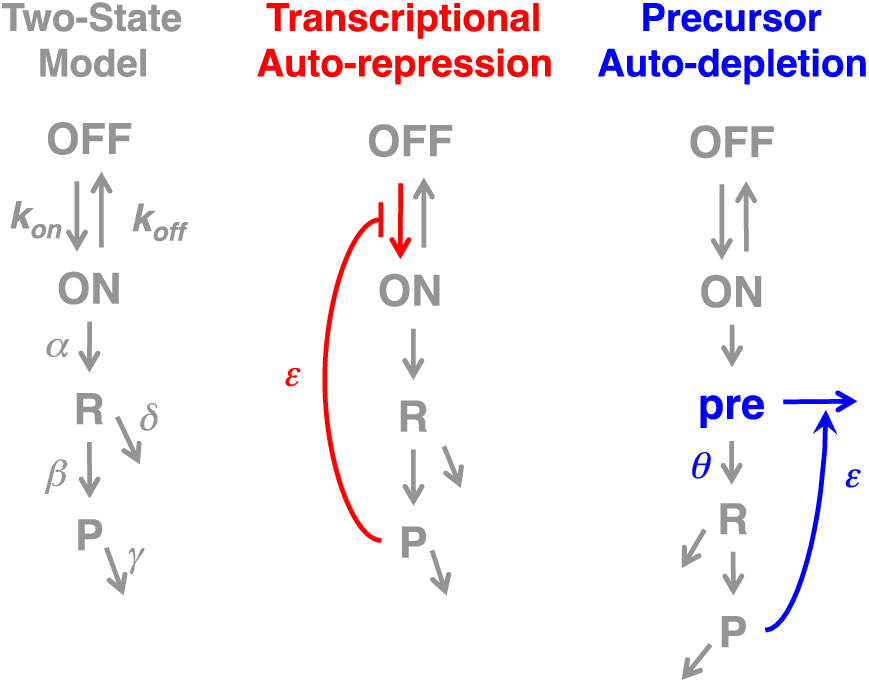

Here, *ON* and *OFF* are active and repressed promoter states, respectively, *R* represents mRNA, *P* represents protein, and all parameters are ‘lumped’ rate parameters representing transcription (*α*), translation (*β*), mRNA degradation (*δ*), protein degradation (γ), and feedback strength (*ε*).

In the auto-depletion model, *pre* is a pre-mRNA that is spliced into *R* at rate *θ*. Steady-state analysis of the deterministic model (Supp. Info) shows negative–feedback strength depends on the relative rates of protein-mediated depletion and splicing (e.g., ε >> *θ* needed for strong feedback).

These models can be numerically simulated (Supp. Info.) to examine the effects of varying feedback (*ε*) on protein levels and noise:

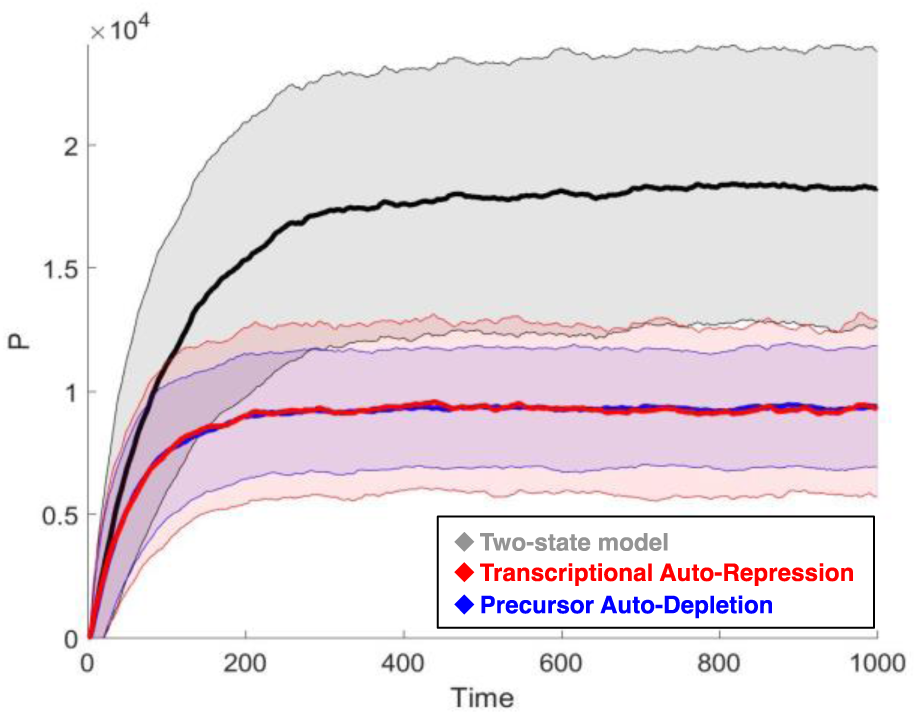

In the two-state model, when the mean is reduced to one half its original value, CV^2^ increases from 0.14 to 0.15 (i.e., Poissonian scaling). Under the same parameters, equivalent reductions in the mean through transcriptional auto-repression reduce CV^2^ to 0.12 whereas precursor auto-depletion reduces the CV^2^ to 0.06 for the same reduction in mean, despite the added noise source (i.e., the extra species, *pre*). In other parameter regimes, auto-repression can actually amplify noise^17,37,42^ whereas auto-depletion avoids this perverse noise amplification (Supp. Info.).

These results agree with analytics for Poissonian (σ^2^ = 〈P〉) expression^42^. For the transcriptional bursting case, analysis^43,44^ shows that σ^2^ = Bb〈P〉 (where *b* is the transcriptional burst size [*α*/*k_off_*] and *B* is the translational efficiency [*β/δ*]), such that the intrinsic noise and mean can reduce to:

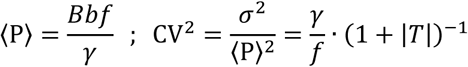

where *f* is the transcriptional burst frequency (1/*k_on_* +1/*k_off_*)^−1^, and *T* is the modification due to feedback strength^44^. Transcriptional auto-repression reduces 〈*P*〉 by decreasing *f* and can thus raise CV^2^ if *T* is not strong enough. In contrast, precursor auto-depletion acts on *B* (and possibly *b*) without perturbing *f*, thus breaking the inverse relation (the added noise source from the *pre* species is high frequency and effectively filtered out at the protein level [Supp. Info.]). Intuitively, this enhanced noise minimization results because precursor auto-depletion acts on RNA, which can have a large number of molecules that can be reduced in an analog-like fashion (from *x* molecules to *x* − *n* molecules). In contrast, transcriptional auto-repression acts on a single-molecule, DNA, which can only toggle between fully active (ON) and repressed (OFF). Thus, transcriptional auto-repression inevitably affects promoter toggling (the major driver of noise) but precursor auto-depletion does not. These analytics generalize and hold beyond two-state models (Supp. Info).

The enhanced SA7 splicing mutants exhibit reduced stability of the active-replication state (Fig. 4, A and B). Destabilization of the active state cannot be explained by the deterministic model (Fig. S13)—the deterministic model in fact predicts that SA7 mutants, which have *increased* Tat concentrations (Fig. S12), would require more time to turn off Tat and establish a latent state. Moreover, establishment of latency appears to occur as an abrupt ‘exchange of mass’ process between bimodal peaks—a hallmark of stochastic transitioning^12^—as opposed to gradual shifting of a single unimodal peak. The accelerated destabilization of the active state in the mutants, despite higher concentrations of Tat, is consistent with the stochastic precursor auto-depletion model in which transitioning is driven by noise rather than mean level of Tat and in which auto-depletion feedback minimizes noise to limit stochastic transitions (Fig. 4C).

**Figure 4:**
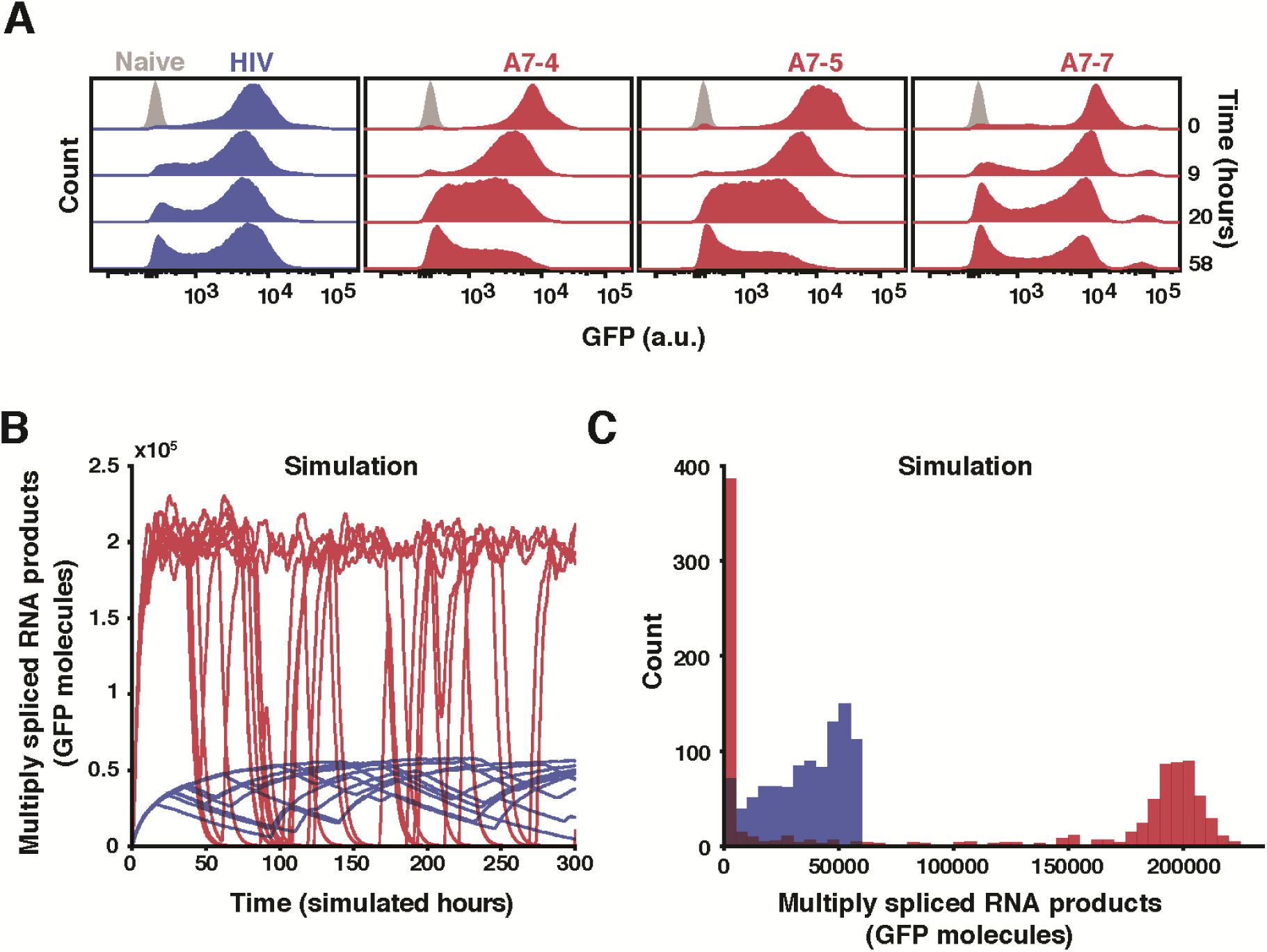
Precursor auto-depletion stabilizes HIV’s active state. (**A**) Flow cytometry analysis of active-state stability following a pulse of TNF reactivation for wild-type HIV d_2_G and SA7 mutants; cells were removed from TNF induction at time 0. (**B**) Representative Gillespie simulations of the HIV precursor auto-depletion model (blue trajectories) showing that GFP-expression noise is attenuated and stochastic ON–OFF switching is minimized. Abrogating precursor auto-depletion (red trajectories) results in increased stochastic ON–OFF switching despite an about 3-fold increase in mean-expression level of the ON state. (**C**) Histograms of 1000 simulations of the HIV precursor auto-depletion model (blue) and abrogated precursor auto-depletion mutant (red) at end of simulation run (i.e., t = 300 hours). In agreement with the flow cytometry data, simulations of the mutant exhibit substantially more trajectories in the GFP OFF state compared to wild type.

Overall, the data indicate that precursor auto-depletion circuitry strongly suppresses transcriptional noise to stabilize HIV active replication after the virus’s noise-driven active-vs.- latent decision. By acting post-transcriptionally, this circuitry overcomes the fundamental theoretical limits on noise suppression by transcriptional feedback^17^; where noise suppression scales with the quartic root of regulator activity (e.g., a decrease in CV to one third of the unregulated level would require an 81-fold change in regulator abundance). In contrast, the precursor auto-depletion circuitry described here shows a one-third reduction in CV^2^ (Fig. 3B), without approaching an 81-fold change in abundance of regulator (Fig. S12).

From an evolutionary perspective, the coupling of noise-amplifying (Tat positive-feedback) circuitry followed downstream by strongly noise-suppressing (Rev auto-depletion) circuitry appears to be expensive regulatory architecture but has the benefit of optimizing fitness for probabilistic bet-hedging strategies^46^. Interestingly, the auto-depletion motif appears to be the equivalent of “parametric control” in electrical signal processing—in which additional parameters are embedded in a circuit to provide more layers of regulation—a concept also exploited by kinetic proofreading in biochemical networks^45^. For HIV, it remains unclear whether this circuitry evolved specifically as an adaptation to suppress noise or is tied to the virus’s need to retain full-length unspliced genomic RNA for packaging. Either way, the circuitry appears conserved^47^—despite alternate non-feedback mechanisms existing for viruses to export unspliced RNA^48^—and its noise buffering effects appear required to stabilize active replication. The conserved nature of this noise buffering circuitry suggests a functional basis for similar regulatory motifs that ‘license’ splicing of precursors^49^. For example, splicing regulation by serene- and arginine-rich (SR) proteins, which shunt their pre-mRNAs to the non-sense mediated decay (NMD) pathway through alternative splicing^50^ may represent a form of licensed auto-depletion. Moreover, the widespread occurrence of post-transcriptionally spliced ‘detained introns’^51^ suggests that auto-depletion circuitry could be an efficient noise attenuation mechanism in diverse systems.

## Author Contributions

L.S.W., W.Y.W., E.I., M.L.S., and B.R. conceived and designed the study. W.Y.W., E.I., and B.R., M.M.K.H., L.S.W. designed and performed the experiments. W.Y.W., C.C., M.L.S., E.I., B.R., R.D.D., M.M.K.H. and L.S.W. analyzed the data and models. M.M.K.H., W.Y.W., E.I., and L.S.W. wrote the paper.

## Acknowledgements

We are grateful to H. Madhani, P. Walter, C. Guthrie, A. Frankel, W. Greene, J. Karn, R. Tsien, A. Hoffmann, J. Guatelli, L. Tsimring, T. Notton, J. Young, and S. Chunda for providing reagents, equipment, and helpful discussions; K. Thorn (Nikon Imaging Center, UCSF) and the UCSF-Gladstone Center for AIDS Research flow core, funded through (P30 AI027763) and (S10 RR028962-01) for invaluable technical expertise. M.M.K.H is supported by an NWO Rubicon fellowship. M.L.S. is supported by the Collective Phenomena in Nanophases Research Theme at the Center for Nanophase Materials Sciences, sponsored at Oak Ridge National Laboratory by the Office of Basic Energy Sciences, U.S. Department of Energy. L.S.W. acknowledges support from the Alfred P. Sloan Research Fellowship, NIH awards R01AI109593, P01AI090935, and the NIH Director’s New Innovator Award (OD006677) and Pioneer Award (OD17181) programs.

## Supplementary Materials

Supplementary Figures S1 to S13

Experimental Materials and Methods

Mathematical Modeling Methods

Supplementary Tables S1 to S12

Supplementary References

Movie S1

## References

1 Acar, M., Mettetal, J. T. & van Oudenaarden, A. Stochastic switching as a survival strategy in fluctuating environments. Nat Genet 40, 471-475, (2008).

2 Balazsi, G., van Oudenaarden, A. & Collins, J. J. Cellular decision making and biological noise: from microbes to mammals. Cell 144, 910-925, (2011).

3 Süel, G. M., Garcia-Ojalvo, J., Liberman, L. M. & Elowitz, M. B. An excitable gene regulatory circuit induces transient cellular differentiation. Nature 440, 545-550, (2006).

4 Süel, G. M., Kulkarni, R. P., Dworkin, J., Garcia-Ojalvo, J. & Elowitz, M. B. Tunability and Noise Dependence in Differentiation Dynamics. Science 315, 1716-1719, (2007).

5 Çağatay, T., Turcotte, M., Elowitz, M. B., Garcia-Ojalvo, J. & Süel, G. M. Architecture-Dependent Noise Discriminates Functionally Analogous Differentiation Circuits. Cell 139, 512-522, (2009).

6 Metzger, B. P., Yuan, D. C., Gruber, J. D., Duveau, F. & Wittkopp, P. J. Selection on noise constrains variation in a eukaryotic promoter. Nature 521, 344-347, (2015).

7 Fraser, H. B., Hirsh, A. E., Giaever, G., Kumm, J. & Eisen, M. B. Noise minimization in eukaryotic gene expression. PLoS Biol 2, e137, (2004).

8 Elowitz, M. B., Levine, A. J., Siggia, E. D. & Swain, P. S. Stochastic gene expression in a single cell. Science 297, 1183-1186, (2002).

9 Alon, U. An introduction to systems biology: design principles of biological circuits. (Chapman & Hall/CRC, 2007).

10 Raser, J. M. & O’Shea, E. K. Control of stochasticity in eukaryotic gene expression. Science 304, 1811-1814, (2004).

11 Blake, W. J., M, K. A., Cantor, C. R. & Collins, J. J. Noise in eukaryotic gene expression. Nature 422, 633-637, (2003).

12 Weinberger, L. S., Burnett, J. C., Toettcher, J. E., Arkin, A. P. & Schaffer, D. V. Stochastic gene expression in a lentiviral positive-feedback loop: HIV-1 Tat fluctuations drive phenotypic diversity. Cell 122, 169-182, (2005).

13 Suel, G. M., Kulkarni, R. P., Dworkin, J., Garcia-Ojalvo, J. & Elowitz, M. B. Tunability and noise dependence in differentiation dynamics. Science 315, 1716-1719, (2007).

14 Chang, H. H., Hemberg, M., Barahona, M., Ingber, D. E. & Huang, S. Transcriptome-wide noise controls lineage choice in mammalian progenitor cells. Nature 453, 544-547, (2008).

15 Cheong, R., Rhee, A., Wang, C. J., Nemenman, I. & Levchenko, A. Information transduction capacity of noisy biochemical signaling networks. Science 334, 354-358, (2011).

16 Schmiedel, J. M. et al. Gene expression. MicroRNA control of protein expression noise. Science 348, 128-132, (2015).

17 Lestas, I., Vinnicombe, G. & Paulsson, J. Fundamental limits on the suppression of molecular fluctuations. Nature 467, 174-178, (2010).

18 Weinberger, L. S. A minimal fate-selection switch. Curr Opin Cell Biol 37, 111-118, (2015).

19 Siliciano, R. F. & Greene, W. C. HIV latency. Cold Spring Harb Perspect Med 1, a007096, (2011).

20 Weinberger, A. D. & Weinberger, L. S. Stochastic Fate Selection in HIV-Infected Patients. Cell 155, 497-499, (2013).

21 Razooky, B. S., Pai, A., Aull, K., Rouzine, I. M. & Weinberger, L. S. A hardwired HIV latency program. Cell 160, 990-1001, (2015).

22 Felber, B. K., Drysdale, C. M. & Pavlakis, G. N. Feedback regulation of human immunodeficiency virus type 1 expression by the Rev protein. Journal of virology 64, 3734-3741, (1990).

23 Becskei, A. & Serrano, L. Engineering stability in gene networks by autoregulation. Nature 405, 590-593, (2000).

24 Fong, Y. W. & Zhou, Q. Stimulatory effect of splicing factors on transcriptional elongation. Nature 414, 929-933, (2001).

25 Taube, R. & Peterlin, M. Lost in transcription: molecular mechanisms that control HIV latency. Viruses 5, 902-927, (2013).

26 Malim, M. H., Hauber, J., Fenrick, R. & Cullen, B. R. Immunodeficiency virus rev trans-activator modulates the expression of the viral regulatory genes. Nature 335, 181-183, (1988).

27 Felber, B. K., Hadzopoulou-Cladaras, M., Cladaras, C., Copeland, T. & Pavlakis, G. N. rev protein of human immunodeficiency virus type 1 affects the stability and transport of the viral mRNA. Proceedings of the National Academy of Sciences of the United States of America 86, 1495-1499, (1989).

28 Ossareh-Nazari, B., Bachelerie, F. & Dargemont, C. Evidence for a role of CRM1 in signal-mediated nuclear protein export. Science 278, 141-144, (1997).

29 Zhang, G., Zapp, M. L., Yan, G. & Green, M. R. Localization of HIV-1 RNA in mammalian nuclei. The Journal of cell biology 135, 9-18, (1996).

30 Vargas, D. Y. et al. Single-molecule imaging of transcriptionally coupled and uncoupled splicing. Cell 147, 1054-1065, (2011).

31 Noble, S. M. & Guthrie, C. Transcriptional pulse-chase analysis reveals a role for a novel snRNP-associated protein in the manufacture of spliceosomal snRNPs. EMBO J 15, 4368-4379, (1996).

32 Waks, Z., Klein, A. M. & Silver, P. A. Cell-to-cell variability of alternative RNA splicing. Mol Syst Biol 7, 506, (2011).

33 Jordan, A., Bisgrove, D. & Verdin, E. HIV reproducibly establishes a latent infection after acute infection of T cells in vitro. EMBO J 22, 1868-1877, (2003).

34 Lahav, G. et al. Dynamics of the p53-Mdm2 feedback loop in individual cells. Nat Genet 36, 147-150, (2004).

35 Hoffmann, A., Levchenko, A., Scott, M. L. & Baltimore, D. The IkappaB-NF-kappaB signaling module: temporal control and selective gene activation. Science 298, 1241-1245, (2002).

36 Weinberger, L. S., Dar, R. D. & Simpson, M. L. Transient-mediated fate determination in a transcriptional circuit of HIV. Nat Genet 40, 466-470, (2008).

37 Austin, D. W. et al. Gene network shaping of inherent noise spectra. Nature 439, 608-611, (2006).

38 Pearson, R. et al. Epigenetic silencing of human immunodeficiency virus (HIV) transcription by formation of restrictive chromatin structures at the viral long terminal repeat drives the progressive entry of HIV into latency. Journal of virology 82, 12291-12303, (2008).

39 Fukuda, M. et al. CRM1 is responsible for intracellular transport mediated by the nuclear export signal. Nature 390, 308-311, (1997).

40 Wu, Y., Beddall, M. H. & Marsh, J. W. Rev-dependent indicator T cell line. Current HIV research 5, 394-402, (2007).

41 Dyhr-Mikkelsen, H. & Kjems, J. Inefficient spliceosome assembly and abnormal branch site selection in splicing of an HIV-1 transcript in vitro. J Biol Chem 270, 24060-24066, (1995).

42 Swain, P. S. Efficient attenuation of stochasticity in gene expression through post-transcriptional control. J Mol Biol 344, 965-976, (2004).

43 Dar, R. D., Razooky, B. S., Weinberger, L. S., Cox, C. D. & Simpson, M. L. The Low Noise Limit in Gene Expression. PLoS One 10, e0140969, (2015).

44 Simpson, M. L., Cox, C. D. & Sayler, G. S. Frequency domain analysis of noise in autoregulated gene circuits. Proceedings of the National Academy of Sciences of the United States of America 100, 4551-4556, (2003).

45 Hopfield, J. J. Kinetic proofreading: a new mechanism for reducing errors in biosynthetic processes requiring high specificity. Proceedings of the National Academy of Sciences of the United States of America 71, 4135-4139, (1974).

46 Rouzine, I. M., Weinberger, A. D. & Weinberger, L. S. An evolutionary role for HIV latency in enhancing viral transmission. Cell 160, 1002-1012, (2015).

47 Hidaka, M., Inoue, J., Yoshida, M. & Seiki, M. Post-transcriptional regulator (rex) of HTLV-1 initiates expression of viral structural proteins but suppresses expression of regulatory proteins. EMBO J 7, 519-523, (1988).

48 Bray, M. et al. A small element from the Mason-Pfizer monkey virus genome makes human immunodeficiency virus type 1 expression and replication Rev-independent. Proceedings of the National Academy of Sciences of the United States of America 91, 1256-1260, (1994).

49 Li, Y. I. et al. RNA splicing is a primary link between genetic variation and disease. Science 352, 600-604, (2016).

50 Lareau, L. F., Inada, M., Green, R. E., Wengrod, J. C. & Brenner, S. E. Unproductive splicing of SR genes associated with highly conserved and ultraconserved DNA elements. Nature 446, 926-929, (2007).

51 Boutz, P. L., Bhutkar, A. & Sharp, P. A. Detained introns are a novel, widespread class of post-transcriptionally spliced introns. Genes Dev 29, 63-80, (2015).

52 Malim, M. H. & Cullen, B. R. HIV-1 structural gene expression requires the binding of multiple Rev monomers to the viral RRE: implications for HIV-1 latency. Cell 65, 241-248, (1991).

